# Development and application of an uncapped mRNA platform

**DOI:** 10.1101/2022.06.05.494796

**Authors:** Xiaodi Zheng, Biao Liu, Peng Ni, Linkang Cai, Xiaotai Shi, Zonghuang Ke, Siqi Zhang, Bing Hu, Binfeng Yang, Yiyan Xu, Wei Long, Zhizheng Fang, Yang Wang, Wen Zhang, Yan. Xu, Zhong Wang, Kai Pan, Kangping Zhou, Hanming Wang, Hui Geng, Han Hu, Binlei Liu

**Affiliations:** National ‘‘111’’ Center for Cellular Regulation and Molecular Pharmaceutics, Key Laboratory of Fermentation Engineering (Ministry of Education), Hubei Provincial Cooperative Innovation Center of Industrial Fermentation, College of Bioengineering, Hubei University of Technology, Wuhan, China; Wuhan Binhui Biopharmaceutical Co., Ltd. Wuhan, China; Hubei Provincial Centre for Disease Control and Prevention, Wuhan, China; Department of Immunology, National Cancer Center/National Clinical Research Center for Cancer/Cancer Hospital, Chinese Academy of Medical Sciences and Peking Union Medical College, Beijing, China; Huazhong Normal University, Wuhan, China

**Keywords:** Uncapped mRNA, LNP, Omicron strain Neutralization

## Abstract

A novel uncapped mRNA platform was developed. Five lipid nanoparticle (LNP)-encapsulated mRNA constructs were made to evaluate several aspects of our platform, including transfection efficiency and durability *in vitro* and *in vivo* and the activation of humoral and cellular immunity in several animal models. The constructs were eGFP-mRNA-LNP (for enhanced green fluorescence mRNA), Fluc-mRNA-LNP (for firefly luciferase mRNA), S^δT^-mRNA-LNP (for Delta strain SARS-CoV-2 spike protein trimer mRNA), gD^ED^-mRNA-LNP (for truncated glycoprotein D mRNA coding ectodomain from herpes simplex virus type 2 (HSV2)) and gD^FR^-mRNA-LNP (for truncated HSV2 glycoprotein D mRNA coding amino acids 1∼400). Quantifiable target protein expression was achieved *in vitro* and *in vivo* with eGFP-and Fluc-mRNA-LNP. S^δT^-mRNA-LNP, gD^ED^-mRNA-LNP and gD^FR^-mRNA-LNP induced both humoral and cellular immune responses comparable to those obtained by previously reported capped mRNA-LNP constructs. Notably, S^δT^-mRNA-LNP elicited neutralizing antibodies in hamsters against the Omicron and Delta strains. Additionally, gD^ED^-mRNA-LNP and gD^FR^-mRNA-LNP induced potent neutralizing antibodies in rabbits and mice. The mRNA constructs with uridine triphosphate (UTP) outperformed those with N1-methylpseudouridine triphosphate (N1mψTP) in the induction of antibodies via S^δT^-mRNA-LNP. Our uncapped, process-simplified, and economical mRNA platform may have broad utility in vaccines and protein replacement drugs.

## 1. Introduction

Messenger RNA (mRNA) is responsible for transcribing the genetic information stored in DNA and directing the synthesis of proteins in cells. Various problems must be addressed to work with mRNA, such as extreme instability, degradation by RNases that occur widely *in vivo* and *in vitro* (Green and Sambrook 2019), dependency on the translation mechanism in the target cells (Eygeris et al. 2022), and the propensity to trigger unnecessary immune responses(Wu et al. 2020). Efforts have been made for decades to obtain mRNA molecules with strong stability, high translation efficiency, and low immunogenicity. The successful development of the new coronavirus mRNA vaccines indicates that mRNA technology has matured and can be used as a powerful, versatile, and fast-response application platform (Anderson et al. 2020).

Studies have found that many naturally existing mRNA chemical modifications have biological significance in mRNA stability. At present, the following chemical modifications are commonly used in mRNA technology: N1- and N6-methyladenosine (m1A, m6A, m6Am) (Davalos et al. 2018), 3- and 5-methylcytosine (m3C, m5C) (Gloge et al. 2014), 5-hydroxymethylcytosine (hm5C) (Fu et al. 2014; Huber et al. 2015), 2’-O-methylation (Nm) (Dai et al. 2017; Dimitrova et al. 2019), and pseudouridine (Ψ) (Davalos et al. 2018; Boo and Kim 2020). Among them, Ψ is the first and most common mRNA modification discovered (Davis and Allen 1957). Katalin Karikó*et al*. have successively found that the introduction of pseudouridine triphosphate (ψTP) into mRNA can reduce its immunogenicity, which decreases with the increase in the proportion of ψTP introduced (Karikó et al. 2005). In 2015, Oliwia Andries *et al*. found that the complete replacement of UTP with N1-methylpseudouridine triphosphate (N1mψTP) reduced the immunogenicity of mRNA and enhanced protein expression more than the complete replacement of UTP with ψTP (Andries et al. 2015). In 2022, Chutamath Sittplangkoon *et al* reported that mRNA vaccine with unmodified uridine could suppress lung metastasis in a melanoma model (Sittplangkoon et al. 2022). The two new coronavirus mRNA vaccines, mRNA-1273 (Moderna, https://www.modernatx.com/research/product-pipeline) and BNT162b2 (Pfizer-BioNTech, https://biontech.de/science/pipeline), both use N1mψTP instead of UTP (Nance and Meier 2021).

To reduce the immunogenicity of mRNA and improve its intracellular stability and translation efficiency, mRNA structures have been designed sequentially to have a 5’ cap, a 5’-untranslated region (UTR), a coding region, a 3’-UTR and a polyadenosine tail (poly A or pA). Cap-dependent structures can use analogs, such as m7GpppG, as a protective structure that shields RNA from exonuclease cleavage and initiates mRNA translation (Sahin et al. 2014). In addition, the internal ribosome entry site (IRES) of some mRNAs has a similar function to the cap structure. In 1988, Pelletier *et al*. found that the 5’-UTR of poliovirus has a P2 sequence of approximately 450 nucleotides that can guide eukaryotic mRNA translation (Pelletier et al. 1988); Jang *et al*. also found a sequence in the 5’-UTR of encephalomyocarditis virus (Jang and Wimmer 1990). In 1991, Macejak *et al*. found that there is an IRES element in the 5’-UTR of the cellular immunoglobulin heavy chain binding protein gene (Macejak and Sarnow 1991). Using a constructed circular mRNA, it was demonstrated that IRES can guide protein synthesis *in vitro*, indicating that protein translation can specifically depend on the IRES sequence (Pyronnet *et al*. 2000). IRES facilitates ribosome assembly and initiates translation by recruiting different trans-acting factors, making it possible for protein translation initiation to occur independently of the 5’ cap structure of the mRNA.

Complementary to a working mRNA technology is a proper delivery technology. The delivery of mRNA to the interior of cells requires addressing enzymatic degradation and membrane barriers caused by electrostatic repulsion. With the approval of two mRNA vaccines, LNPs are recognized as the most successful mRNA delivery vehicles (Teo 2022). Liposomes are vesicle structures composed of lipid molecules that were discovered by scientists A. D. Bangham and R. W. Horne under a microscope as early as 1961(Bangham et al. 1962; Bangham and Horne 1964). In 1995, the first liposomal drug Doxil approved by the FDA in the US was used for ovarian cancer and breast cancer chemotherapy through the HSPC/DMG-PEG dual component liposome encapsulated drug doxorubicin to reduce the toxicity of free drugs to other organs (Barenholz 2012). For the mRNA vaccine against the new COVID-19 coronavirus disease, Moderna used an ionizable lipid SM-102 (Cho et al. 2021), while Pfizer and BioNTech used an ionizable lipid called ALC-0315 (Crommelin et al. 2021). LNPs can not only effectively deliver mRNA to specific target cells to prevent mRNA from being degraded or cleared but also help mRNA escape from cellular endosomes to the cytoplasm in a timely manner and be translated into corresponding proteins (Maugeri et al. 2019).

Given the rapid development of mRNA technology and the urgent application needs, the mRNA platform described in this study is composed of an uncapped mRNA structure without using N1mψTP in place of UTP. Our mRNA platform aims to improve product manufacturing efficiency, reduce costs, improve efficacy and ensure quality. By further optimizing the mRNA process and technical route, it is expected that our platform can provide more effective, longer-lasting and more efficient diversified treatment options for infectious diseases, rare diseases, cancer, chronic diseases and other diseases.

## 2. Materials and methods

### 2.1. Plasmid construction

All the constructed plasmids are derived from pT7AMP (constructed in our laboratory, to be published soon). The coding sequences for two truncated D-type envelope glycoproteins from herpes simplex virus type 2 (HSV2 gD^ED^ with amino acid sequence 1-316 and HSV2 gD^FR^ with amino acid sequence 1-400), the coding sequence for SARS-CoV-2 spike protein trimer from Delta strain, termed S^δT^ with amino acid sequence 1-1274aa (including proline mutations at positions 982 and 983, RRAR to GGSG mutations at 678-681aa for S1/S2 cleavage site and a trimerization domain at positions 1203-1274aa), the coding sequence for firefly luciferase (Fluc), the coding sequence for enhanced green fluorescent protein (eGFP) and the sequence for poly A (>100 bp) were synthesized by Nanjing GenScript Biotechnology Co., Ltd. (Nanjing, China) (Chaudhary et al. 2021). The above synthesized fragments were cloned into the *BglII* site of the ampicillin-resistant pT7AMP vector by homologous recombination to obtain plasmids pT7AMP-gD^ED^, pT7AMP-gD^FR^, pT7AMP-S^δT^, pT7AMP-Fluc, and pT7AMP-eGFP. All the constructed plasmids were verified by sequencing.

### 2.2. In vitro transcription (IVT) and mRNA purification

The plasmids described above were used as templates and primer pairs (T7P-F: agggaataagggcgacacggaaatgttgaatactcat and pA-R: ggattgggaagacaatagcaggcatgctgggg, synthesized by Wuhan Qingke Biotechnology Co., Ltd.) were used for PCR amplification with the high-fidelity enzyme 2×Phanta^®^ Flash Master Mix (Vazyme, Nanjing, China) to obtain mRNA transcription templates. A small amount of PCR product was taken for concentration detection using a Qubit 4.0 Fluorometer (Thermo Fisher Scientific, Waltham, MA, USA). The PCR product quality was also assayed by electrophoresis on a 1% agarose gel at 120 V for 30 minutes followed by visualization with a gel imaging system.

IVT was optimized by DOE (Design Of Experiment). The optimized reaction system contained: ATP 5 mM, CTP 5 mM, GTP 5 mM, UTP 5 mM, recombinant T7 RNA polymerase 1200U, DNA template (20-35 ng/μL), inorganic pyrophosphatase 3U (Roche, Mannheim, Germany), RNase inhibitor 40U (Roche, Mannheim, Germany), 10×Transcription Buffer 20 µL. To synthesize N1-methyl pseudouridine triphosphate (N1mψTP)-modified mRNA, UTP was replaced with N1mψTP (Wuhan Tangzhi Biotechnology Co., Ltd., Wuhan, China). After 3 h at 37 °C, DNase I was added to degrade the DNA in the reaction system.

mRNA was purified using an AKTA Avant chromatography system (GE, Sweden) and a CIMmultus™ Oligo dT 18 chromatography column (BIA Separations, Slovenia). The mRNA sample was mixed with OA (Oligo dT Adjust) buffer at a ratio of 4:1. The mixed sample was injected into the AKTA sample loop. Then, the column was washed with OW (Oligo dT Wash) buffer. The mRNA was eluted from the column using water for injection (in-house). Finally, the concentration and purity of the mRNA stock solution were detected by SEC-HPLC (SHIMADZU, Japan) using an analytical TSK G6000 PWXL column (TOSOH, Japan).

### 2.3. Ionizable lipid synthesis

Lipid-VI was synthesized via heptadecan-9-yl 8-((2-hydroxyethyl)amino)octanoate, (9Z,12Z)-octadeca-9,12-dien-1-yl 6-bromohexanoate and N,N-Diisopropylethylamine in ethyl alcohol at 65°C. Forty-eight hours later, the solvent was removed by rotary evaporation, and crude Lipid-VI was purified by silica gel column chromatography with elution of dichloromethane: methanol = 10:1 (v/v) to obtain Lipid-VI. The construction of Lipid-VI was confirmed using ^1^H nuclear magnetic resonance. (^1^H NMR (400 MHz, CDCl_3_) δ 0.82 – 0.94 (t, *J* = 6.2 Hz, 9H), 1.16 – 1.40 (m, 50H), 1.44 – 1.53 (m, 7H), 1.56 – 1.69 (dd, *J* = 4.8, 11.1 Hz, 6H), 2.00 – 2.10 (q, *J* = 6.8 Hz, 4H), 2.24 – 2.35 (dt, *J* = 7.5, 9.3 Hz, 4H), 2.43 – 2.52 (dt, *J* = 5.1, 8.1 Hz, 4H), 2.57 – 2.66 (t, *J* = 5.3 Hz, 2H), 2.72 – 2.83 (t, *J* = 6.4 Hz, 2H), 3.49 – 3.59 (t, *J* = 5.3 Hz, 2H), 4.01 – 4.09 (t, *J* = 6.8 Hz, 2H), 4.82 – 4.91 (m, 1H), 5.31 – 5.42 (tt, *J* = 6.2, 11.3 Hz, 4H))

### 2.4. Preparation and Characterization of lipid nanoparticles (LNPs)

The LNP formulation was optimized on the basis of the literature (Pallesen et al. 2017). Four components, including SM102 (Sinopeg, Xiamen, China), or our self-designed ionizable amino lipids Lipid-VI, 1,2-distearoyl-sn-glycero-3-phosphocholine (DSPC), cholesterol and 1,2-dimyristoyl-rac-glycerol-3-methoxy polyethylene glycol-2000 (DMG-PEG2000) (AVT, shanghai, China), were dissolved in ethanol at a molar ratio of 50:10:38:2. The obtained lipid mixture was then mixed with 25 mM sodium acetate (pH=5.0) buffer at 1:4 (ethanol:water) in a microfluidic mixer (Precision Nanosystems, Canada or NanoPro System, self-developed). The ionizable N:P ratio in the formulation was 4:1. The encapsulated mRNA-LNPs were concentrated by changing the buffer using a centrifugal filter (Amicon Ultra, Millipore, 30 kDa) or a Tangential Flow Filtration (TFF) membrane (Sartorius, 30 kDa). The mRNA-LNP concentrates were sterilized through a 0.22-micron filter. Then use Malvern particle size instrument Zetasizer Pro (Zetasizer Pro, Malvern, UK) to determine its particle size

### 2.5. Detection of mRNA encapsulation efficiency and In vitro transfection

mRNA concentrations were measured on a Qubit 4.0 Fluorometer (Thermo Fisher Scientific, Waltham, MA, USA) using the Qubit® RNA BR Assay kit. According to the instructions, the unencapsulated mRNA content (C_f_) and the mRNA content (C_t_) of the same sample after adding 0.1% (v/v) Triton X-100 (Sigma) were measured to calculate the mRNA encapsulation efficiency. The encapsulation rate was calculated as (1-C_f_/C_t_)×100%.

BHK cells were seeded in 6-well plates at 2×10^5^ cells per well. After 24 h, the medium in the well was replaced with 2 mL of fresh high-glucose DMEM containing 5% FBS. eGFP-mRNA-LNP (at 3-4 μg mRNA per well) was slowly added along the wall of the well, and the plate was shaken evenly. The cells were cultured for 24 h, followed by microscopic observation of the cells transfected with eGFP-mRNA-LNP.

### 2.6. Animal experiments

*In vivo* transfection of Fluc-mRNA-LNP: LNP-encapsulated Fluc-mRNA was injected intramuscularly in the thighs of mice. At designated time points, 100 µL of D-fluorescein potassium salt was intraperitoneally injected, and images were recorded using an IVIS imaging system (Perkin Elmer, Waltham, MA, USA).

Immunization of S^δT^-mRNA-LNP: At two time points (mice at days 0 and 14; hamsters at days 0 and 21), LNP-encapsulated S^δT^ mRNA was injected into the thigh muscle of mice or subcutaneously in hamsters. Immunization with HSV2 gD-mRNA-LNP: At three time points (days 0, 14, and 49), LNP-encapsulated gD-mRNA products were injected intramuscularly into the thighs of mice. At two time points (day 0 and day 14), different doses of LNP-encapsulated gD-mRNA preparations were injected into the rabbits at the posterior neck intradermally or at the hind leg intramuscularly, respectively. Serum samples were separated for neutralizing antibody detection.

ELISA detection of antibodies: The microtiter plate was coated with SARS-CoV-2 (B.1.617.2) S protein (Vazyme, Nanjing, China) at 100 ng per well and left overnight at 4 °C. The coated plates were washed with wash buffer and blocked with 1% BSA at 37 °C for 2 h. Mouse serum samples were serially diluted 2-fold starting from 1:1000. The hamster serum samples were diluted to 1:2×10^4^ and then serially diluted 2 times. Serially diluted serum samples were added to ELISA plates and incubated at 37 °C for 1 h. The plates were washed and incubated with (HRP)-conjugated antibody that were diluted in wash buffer containing 0.2% BSA. Plates were incubated for 1 h at 37 °C. TMB was added and the reaction was stopped with 2 M sulfuric acids, and the absorbance was measured at 450 nm using a microplate reader.

Detection of neutralization antibodies: Delta (YJ20210701-01) and Omicron (249099) variants of SARS-CoV-2 were provided by Hubei Provincial Center for Disease Control and Prevention. All SARS-CoV-2 live virus experiments were performed in the Biosafety Level 3 (BSL-3) facility at the Institute of Health Inspection and Testing, Hubei Provincial Center for Disease Control and Prevention.

Neutralization of serum antibodies was assessed by a constant virus amount. Serum samples were serially diluted 2-fold starting from 1:8. Each of the diluted serum samples was thoroughly mixed with the oHSV2-eGFP (an HSV2 virus expressing eGFP made in our lab) virus or SARS-CoV-2 (Delta and Omicron strains) solution containing 100 CCID_50_ (or TCID_50_) to obtain a serum-virus mixture that was incubated at 37 °C for 1 h. Then, each serum-virus mixture was inoculated into the corresponding well of a 96-well microtiter plate cultured with a monolayer of Vero cells and incubated for 48-96 h. Fluorescence expression for gD neutralization antibody titer or cytopathic effect (CPE) for SARS-CoV-2 S protein neutralization antibody titer in the wells was observed using the High Content Analysis System (Perkin Elmer, Waltham, MA, USA) or microscope, respectively. Antibody neutralization titers were assessed by the Reed-Muench method.

Alternatively, the neutralizing activity of serum antibodies was assessed by constant serum dilutions. The oHSV2-eGFP virus was serially diluted 2 times from 1:2 to obtain suspensions containing different amounts of virus (1500-100000 CCID_50_). Each virus suspension was thoroughly mixed with serum to obtain a virus-serum mixture that was incubated at 37 °C for 1 hour. Then, each virus-serum mixture was inoculated into the corresponding well of a 96-well microtiter plate cultured with a monolayer of Vero cells and incubated for 48 h. Fluorescence expression in wells (representing cytopathic effect, CPE) was detected using the High Content Analysis System. The neutralizing activity of serum antibodies was evaluated by CPE.

ELISpot tests on animals immunized with uncapped mRNA-LNP: At 2-3 time points (days 0, 14, and 21), LNP-encapsulated mRNA was injected intramuscularly into the thighs of mice. At a designated time point, the immunized mouse spleens were removed, and splenocytes were isolated for the detection of specific T cells.

The detection steps of gD-specific or S^δT^-specific T cells were as follows: 3-5×10^5^ splenocytes per well and 10^6^ CCID_50_ per well OH2 virus inactivated by UV for 30 minutes (or 4 μg per well SARS-CoV-2 S Protein) were mixed and incubated for 48. And then the plates were incubated with biotinylated anti-IFN-γ antibody and streptavidin-ALP for 2 h after washing with PBS. IFN-γ ELISpot was measured by an ELISpot reader (Autoimmun Diagnostika GmbH, Strassberg, Germany). Student’s *t-*test was performed in GraphPad Prism v8.0.1 software to statistically analyze the data.

## 3. Results

### 3.1. Amplification of IVT template and preparation of uncapped mRNA with high yield and purity

As shown in Figure 1A, the element sequences of the mRNA structure include the T7 promoter (T7P), internal ribosome entry sites (IRES), gene of interest (GOI) and poly A. Five GOIs used in this study (gD^ED^, gD^FR^, S^δT^, Fluc or eGFP) were each inserted between the IRES and poly A sequences of pT7AMP by homologous recombination. All constructed plasmids were verified by sequencing.

**Figure 1.**
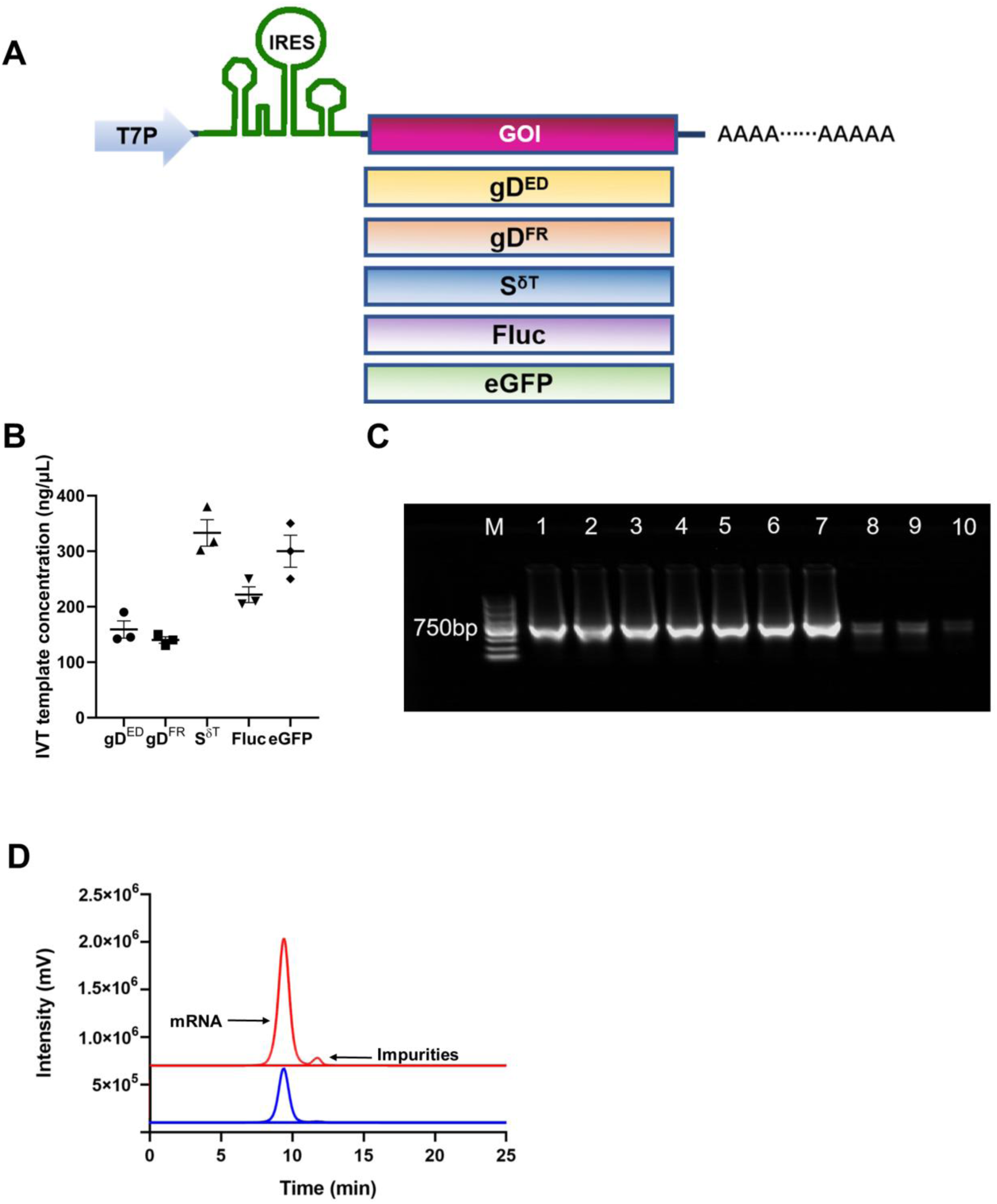
Amplification of the IVT template and preparation of uncapped mRNA with high yield and purity. **A** Schematic diagram of mRNA structures. T7P and pA denote the T7 promoter sequence and polyadenosine sequence, respectively. **B** Scatter plot of IVT template concentrations obtained by PCR amplification from 5 plasmid templates. Data are presented as the mean ±SEM (n = 3). **C** Agarose gel (1%) electropherogram of IVT-synthesized mRNA. Lanes 1∼7 were the 7-repeat IVT products, the sample loading volume was 1 μL, and all IVT templates were Fluc mRNA; M was a 5000 bp DNA marker with a loading volume of 5 μL; lanes 8, 9 and 10 were 900, 500, and 250 ng RNA references, respectively. **D** SEC-HPLC analysis of mRNA samples obtained before and after purification. Red and blue diagrams represent pre- and post-purified mRNA samples. The major peak area of the purified sample was greater than 99%.

The IVT templates were obtained by PCR amplification using the above five plasmids as templates and primer pairs (T7P-F vs. pA-R). The electrophoresis results showed that the five IVT template fragments were all single bands with strong brightness, which was consistent with the expected fragment sizes. The average concentration of IVT templates was 240 ±100 ng μL^-1^, as determined by the Qubit method (Figure 1B). Repeated experiments showed that PCR amplification could rapidly obtain IVT templates with relatively high concentration and purity.

### 3.2. IVT was performed with a PCR-amplified IVT template under DOE-optimized conditions

The mRNA purity was assayed by electrophoresis, and the concentration was determined by electrophoresis and the Qubit assay, and both of these measurements were monitored posttranscriptionally. The concentrations of mRNA synthesized by 7 repeated IVT experiments were all within 4.3 ± 0.5 mg mL^-1^, indicating that the mRNA yield synthesized by IVT optimized by DOE was high and robust (Figure 1C). The mRNA synthesized by IVT was purified by a CIMmultus^TM^ Oligo dT 18 chromatographic column, and a small amount of the purified mRNA was tested for its purity and concentration by HPLC. The SEC-HPLC test results showed that the number and area of cluttered peaks in the purified mRNA samples were reduced, indicating that the mRNA single peak area accounted for 99% of the total peak area (Figure 1D). ***Optimized encapsulation process can stably produce high-quality mRNA-LNPs*.**

A total of six uncapped mRNA constructs, including gD^ED^-mRNA, gD^FR^-mRNA, S protein mRNA from the Delta strain (S^δT^-mRNA), S^δT^-mRNA^N1mψTP^ (N1mψTP replaced UTP), Fluc-mRNA, and Fluc-mRNA^N1mψTP^ were encapsulated. The mRNA and lipid were mixed in a microfluidic mixer (Figure 2B) to produce LNPs, which were analyzed by 1% agarose gel electrophoresis. After electrophoresis, all encapsulated mRNAs could not be removed and remained in the loading wells, and mRNA bands of expected sizes were not seen in the corresponding lanes (Figure 2C). The results of mRNA concentration measurements showed that the encapsulation efficiency of the six mRNA molecules was greater than 90%. Table 1 shows the free mRNA content (C_f_), total mRNA content (C_t_) and encapsulation efficiency values after encapsulation of the six mRNA molecules. After removal of the nonaqueous solvent in the encapsulation system, each mRNA-LNP had a particle size (Z average) between 70 and 100 nm and a polymer dispensability index (PDI) less than 0.2 (Figure 2D, Table 1).

**Figure 2.**
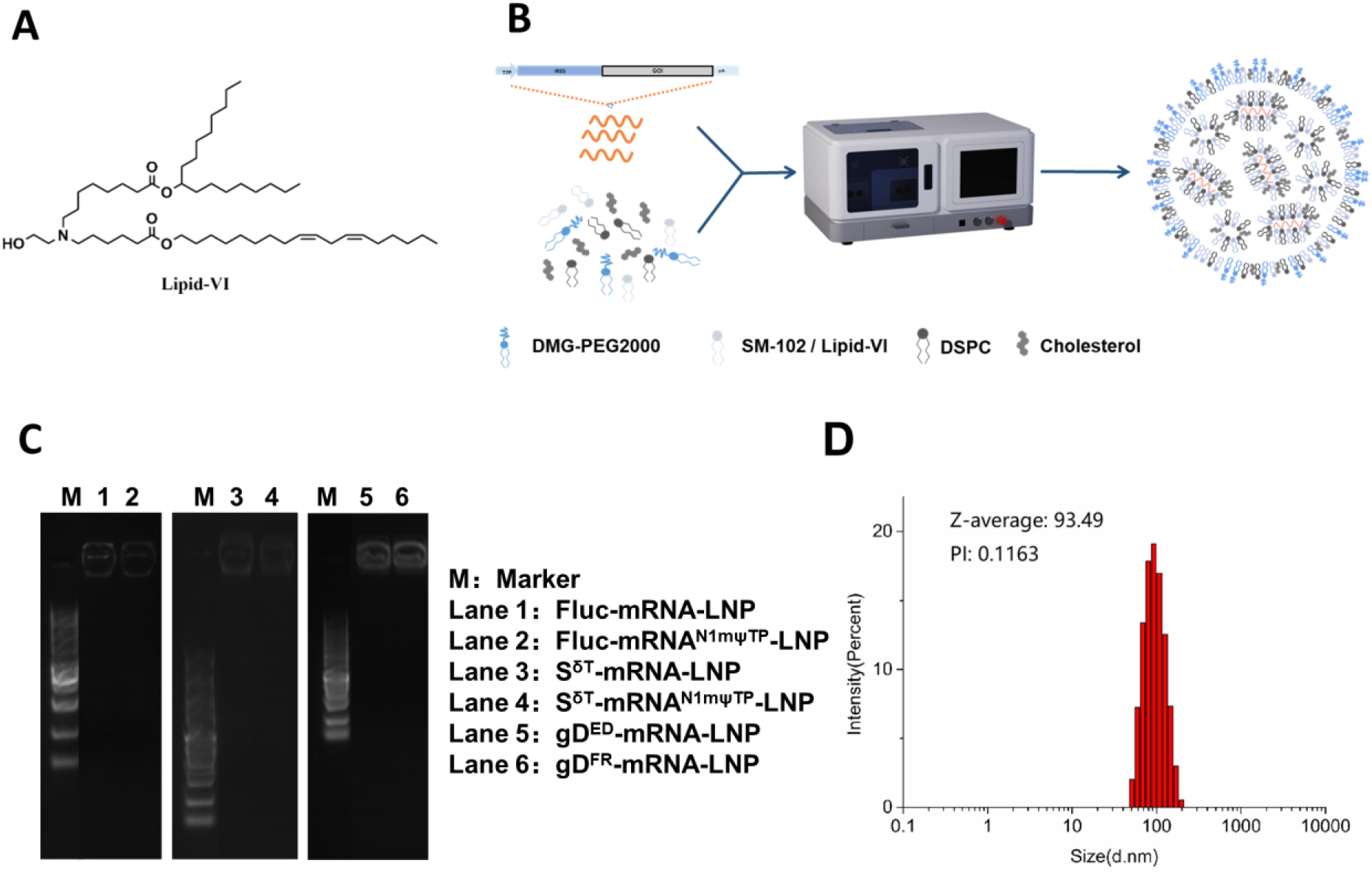
Encapsulation of mRNA. A The structure of Lipid-VI. **B** Process flow diagram. **C** mRNA-LNP electrophoresis with parameters of 1% agarose gel, 120 V and TAE running buffer. M: Marker; Lane 1: Fluc-mRNA-LNP; Lane 2: Fluc^N1mψTP^-mRNA-LNP; Lane 3: S^δT^-mRNA-LNP; Lane 4: S^δT^-mRNA^N1mψTP^-LNP; Lane 5: gD^ED^-mRNA-LNP; Lane 6: gD^FR^-mRNA-LNP. **D** Typical particle size of mRNA-LNP detected with a Mastersizer Laser Diffraction Particle Size Analyzer.

**Table 1.** Test data of key indicators of six mRNA-LNPs.

| | $C_f$ (ng $\mu\text{L}^{-1}$) | $C_t$ (ng $\mu\text{L}^{-1}$) | EN (%) | Z-average (nm) | PDI |
| --- | --- | --- | --- | --- | --- |
| Fluc-mRNA-LNP | 7.18 | 217 | 96.3 | 79.9 | 0.09 |
| Fluc-mRNA<sup>N1mψTP</sup>-LNP | 5.44 | 258 | 97.9 | 74.5 | 0.12 |
| S<sup>δT</sup>-mRNA-LNP | 5.54 | 290 | 98.1 | 80.9 | 0.07 |
| S<sup>δT</sup>-mRNA<sup>N1mψTP</sup>-LNP | 5.40 | 300 | 98.2 | 84.1 | 0.06 |
| gD<sup>ED</sup>-mRNA-LNP | 11.0 | 320 | 96.5 | 85.6 | 0.08 |
| gD<sup>FR</sup>-mRNA-LNP | 12.0 | 265 | 95.4 | 86.7 | 0.07 |

### 3.3. In vitro and in vivo expression of mRNA-LNPs containing indicator genes

The luciferase-fluorescence intensities and durability kinetics of our uncapped mRNA structure and the capped N1mψTP-mRNA structure were detected and found that they were basically comparable (Figure 3A and B). Furthermore, lower doses of uncapped mRNA structure were tested on the animal model. The results also showed that the luciferase-fluorescence intensities of the uncapped mRNA structure and the capped N1mψTP-mRNA structure were similar while the capped UTP group exhibited the weakest signal (Figure 3C and D). These results indicated that the encapsulated Fluc-mRNA uncapped structure can be expressed efficiently *in vivo*.

**Figure 3.**
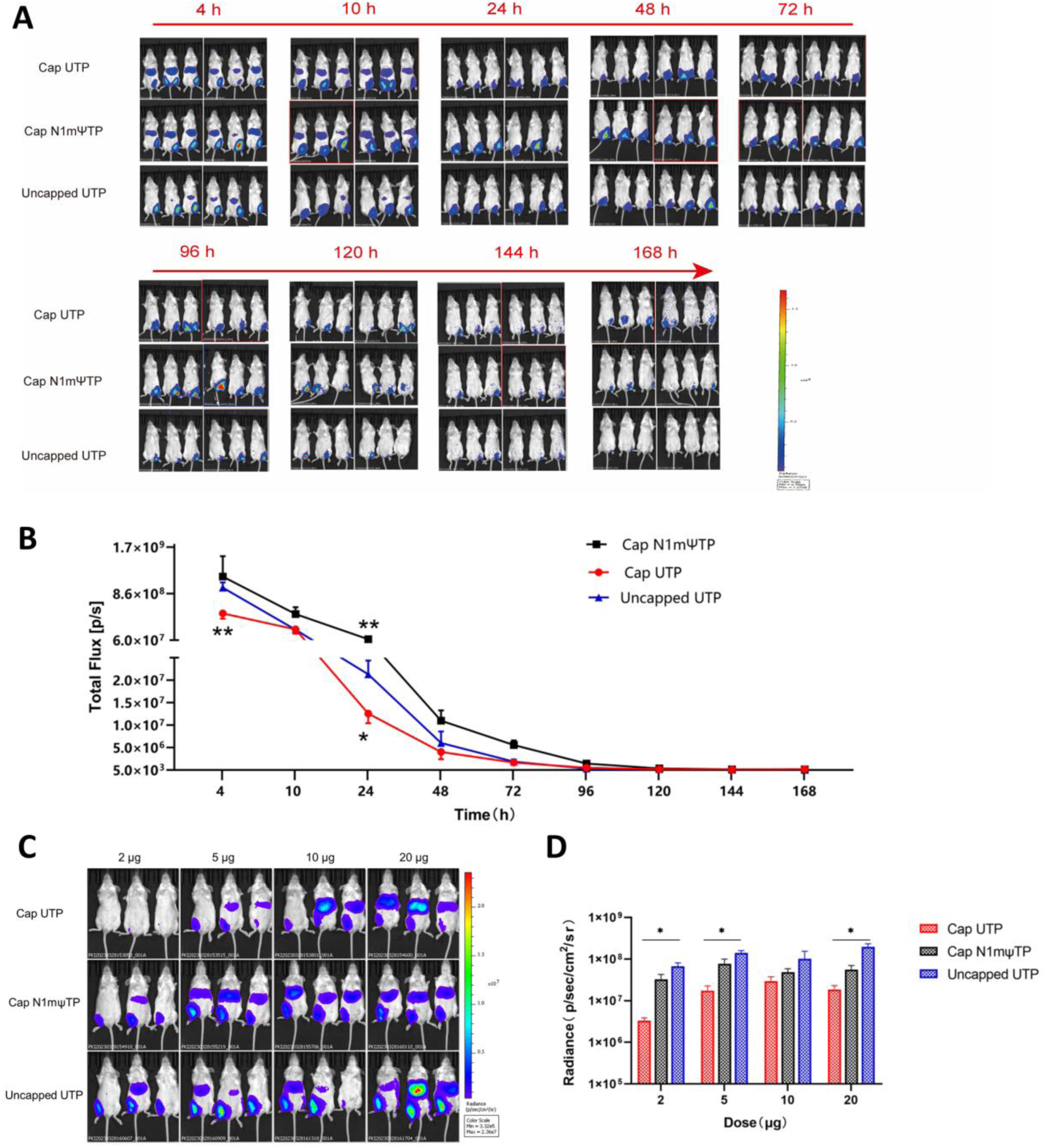
Uncapped mRNA-LNP transfection *in vivo*. **A** The images of fluorescence intensity and **B** the fluorescence quantitative line chart at various time points of the uncapped mRNA, the capped N1mψTP-mRNA, and the capped UTP-mRNA. **C** The images of fluorescence intensity and **D** the fluorescence quantitative histogram at 4 hours post-transfection of the uncapped mRNA, the capped N1mψTP-mRNA, and the capped UTP-mRNA. Significance was calculated using Student’s *t-*test (ns, not significant; *p < 0.05; **p < 0.01; ***p < 0.001).

Our uncapped eGFP-mRNA encapsulated with SM102, Lipo 3000 or self-developed Lipid-VI could easily and efficiently transfect cells *in vitro* to express eGFP (Figure 4A and B). After 24 hours of transfection with eGFP-mRNA-LNP, eGFP could be observed in BHK, indicating that the eGFP-mRNA-LNP can quickly enter the cytoplasm through endocytosis and efficiently be translated. Morphology and boundary of transfected cells was clear, showing that there was no obvious cytotoxicity of the eGFP-mRNA-LNP. The SM102 and Lipid-VI groups all showed good transfection efficiency with the GFP positive rates being above 95% (Figure 4B). Furthermore, *in vivo* assay was performed to compare SM102 and Lipid-VI. The luciferase-fluorescence intensities and durability kinetics of SM102 group and Lipid-VI group were similar (Figure 4C and D). It was further confirmed that both SM102-encapsulated and Lipid-VI-encapsulated S^δT^-mRNA-LNP produced high-titer IgG antibodies when used to immunize Syrian hamsters (Figure 4E and F).

**Figure 4.**
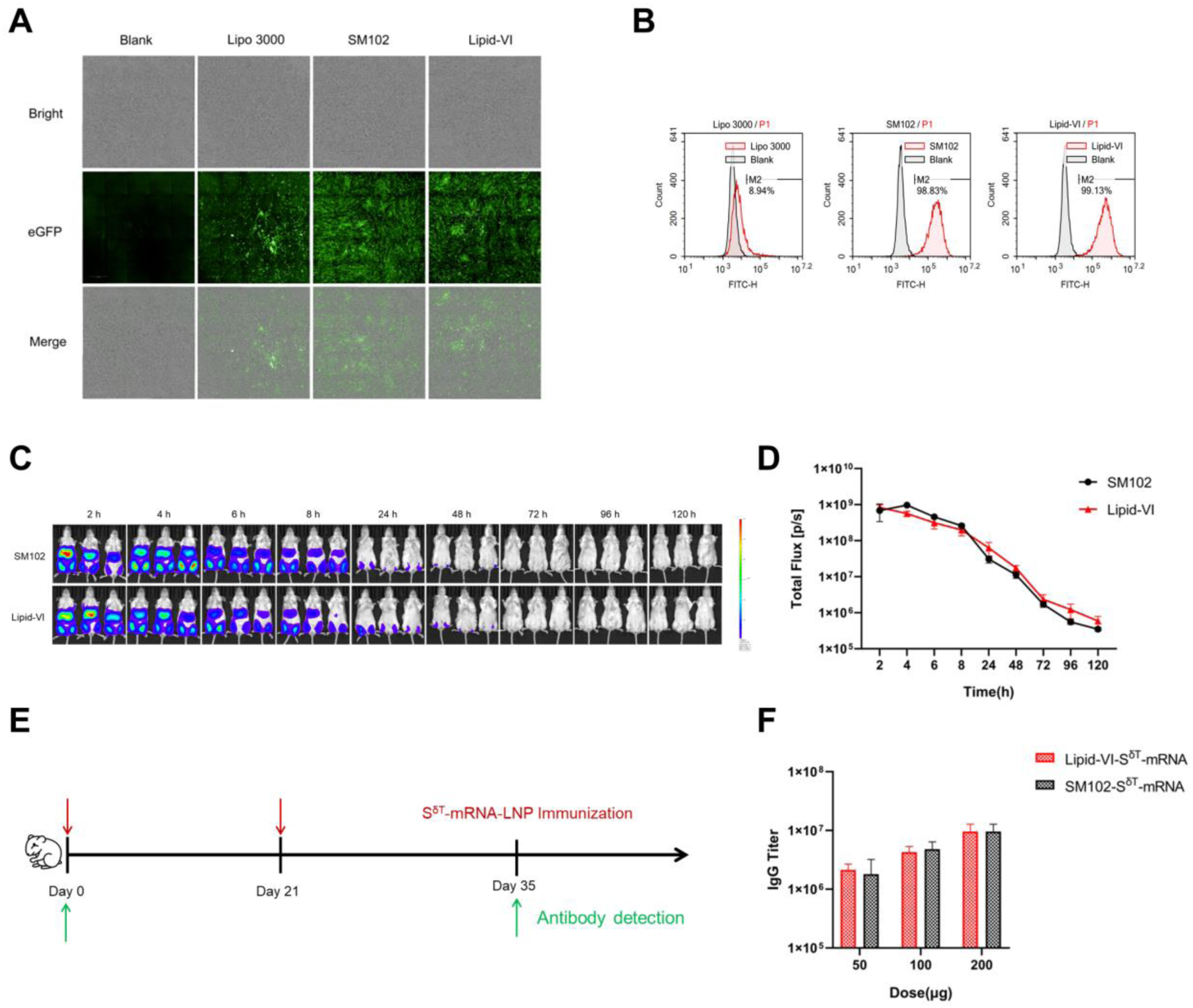
*In vitro* and *in vivo* expression of eGFP-mRNA encapsulated with SM102 and Lipid-VI. **A** Bright-field and dark-field photos of BHK cells transfected with eGFP-mRNA-LNP for 24 hours and **B** the transfection efficiency was detected by flow cytometry. **C** The images of fluorescence intensity and **D** the fluorescence quantitative line chart at various time points post-transfection of SM102 and Lipid-VI encapsulated Fluc-mRNA (30 μg per animal) into mouse muscle. **E** Schematic diagram of immunization and sample collection in Syrian hamsters. **F** The SARS-CoV-2-specific IgG antibody titers of the S^δT^-mRNA encapsulated with SM102 and Lipid-VI on day 35 were determined by ELISA.

### 3.4. Animals immunized with S^δT^-mRNA-LNP but not S^δT^-mRNA^N1mψTP^-LNP produced high-titer binding and potent neutralizing antibodies

Day 0 was defined as the day of the first immunization in experiment 1, and mice were immunized on days 0 and 14 (Figure 5A). The experiment was divided into three groups (n = 3 per group): the UTP group; the N1mψTP group; and the naïve group. On the 28th day, blood was collected to separate the serum and detect the serum IgG antibody titer. The results showed that the average titer of the UTP group reached 26 666, whereas the titers for the N1mψTP group and the naïve group were under the detection limit (Figure 5B). N1mψTP might change the structure of mRNA, causing low expression of S^δT^. The titer of the UTP group was significantly higher than that of the naïve group and the N1mψTPU group.

**Figure 5.**
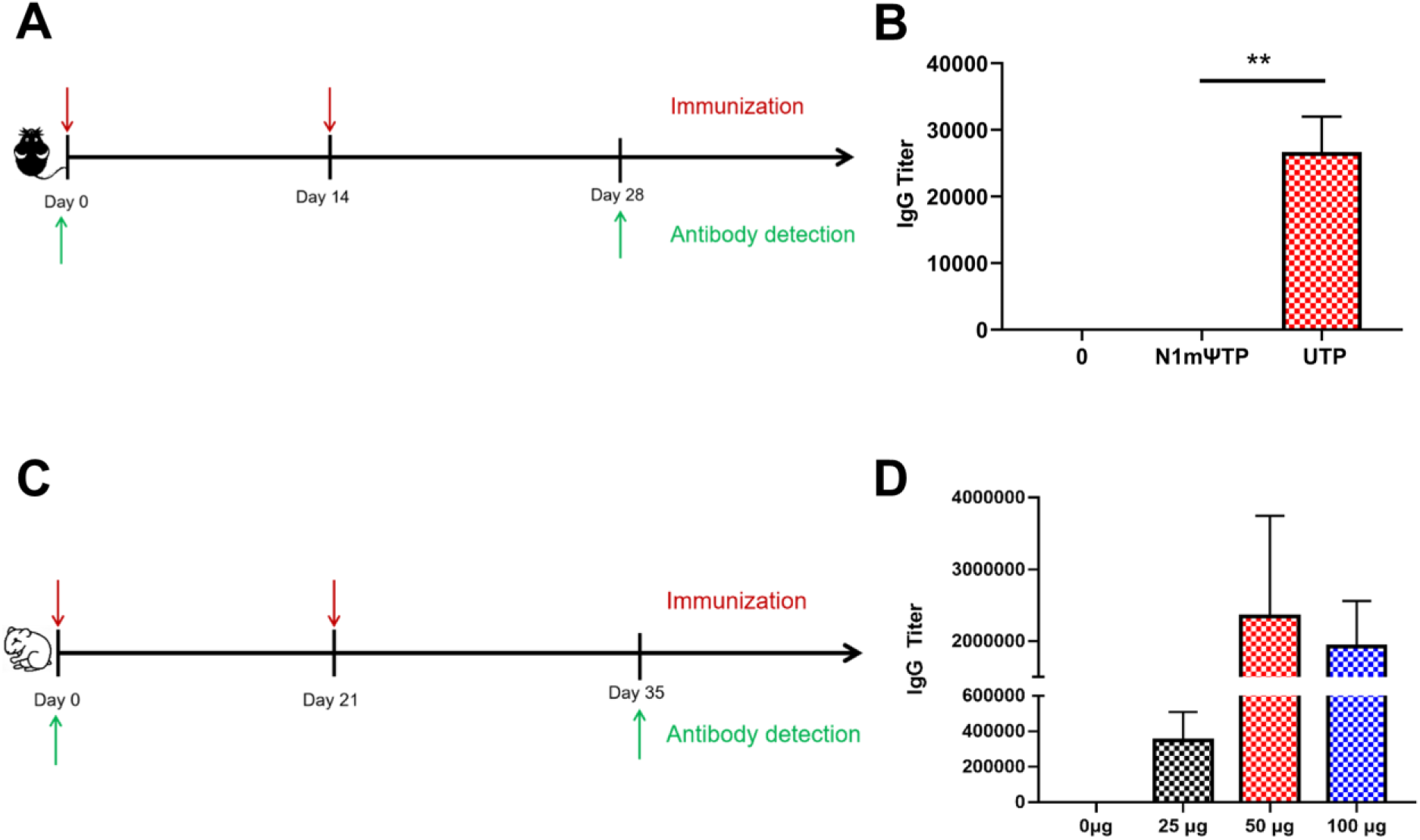
Antibody detection after immunization with S^δT^-mRNA-LNP and S^δT^-mRNA^N1mψTP^- LNP. **A** Schematic diagram of immunization and serum sample collection in C57BL/6 mice. **B** The SARS-CoV-2-specific IgG antibody titers of the 3 groups (0 µg, 30 μg N1mψTP, and 30 μg UTP) were determined by ELISA. Significance was calculated using Student’s *t-*test (ns, not significant; *p < 0.05; **p < 0.01; ***p < 0.001). **C** Schematic diagram of immunization and sample collection in Syrian hamsters. **D** The SARS-CoV-2-specific IgG antibody titers of the 4 groups on day 35 were determined by ELISA.

In experiment 2, the day for the first dose was designated as day 0, and Syrian hamsters were immunized with an intramuscular injection of S^δT^-mRNA-LNP on days 0 and 21 (Figure 5C). The experiment contained four groups (n = 3 per group): the UTP groups of different doses; and the naïve group. On the 35th day, blood samples were collected to separate sera and detect serum binding (IgG) and neutralizing antibody titers. As shown in Figure 5D, the average titers of IgG antibodies in the three experimental groups were 3.47×10^6^ ±1.62×10^6^, 2.35×10^7^ ± 1.40×10^7^, and 1.92×10^7^ ± 0.64×10^7^, respectively. It was noted that the antibody titers in experiment 2 were significantly higher than those in experiment 1, possibly due to differences in animal species, immunization doses and dose regimens. The antibodies raised in Syrian hamsters with S^δT^-mRNA-LNP can effectively neutralize both Delta and Omicron strains of SARS-CoV-2. The S^δT^-mRNA-LNP doses and neutralization titers against Delta and Omicron strains are shown in Figure S1, which indicated that the S^δT^-mRNA-LNP was able to induce potent neutralization antibodies to both Delta and Omicron strains after two immunizations.

### 3.5. Animals immunized with gD^ED^-mRNA-LNP and gD^FR^-mRNA-LNP produced potent neutralizing antibodies

In the mouse experiment, the first immunization was designated as day 0. On days 0, 14, and 49, mice in the experimental groups were injected intramuscularly with gD^ED^-mRNA-LNP (30 μg per mouse, n = 5) or gD^FR^-mRNA-LNP (30 μg per mouse, n = 5) (Figure 6A). On days 0, 28, and 56, blood samples were collected to separate sera for the detection of neutralizing antibodies. One microliter each of serum and oHSV2-eGFP virus containing 6250 CCID_50_ were mixed well and incubated for 1 hour and then inoculated into the well containing a monolayer of Vero cells, and the neutralizing titer of the serum antibody was assessed by eGFP expression and plaque formation. As shown in Figure 6B, all cells treated with the serum-virus mixture in the unimmunized group produced eGFP, and typical viral plaques resulting from virus replication could be seen. In contrast, eGFP and viral plaques were not observed in cells treated with the serum-virus mixture (containing 6250 CCID_50_ virus) in the gD^ED^-mRNA-LNP and gD^FR^-mRNA-LNP groups. The results indicated that the two HSV2 gD antigens expressed *in vivo* from our uncapped mRNA constructs could induce neutralizing antibodies in immunized C57BL/6 mice.

**Figure 6.**
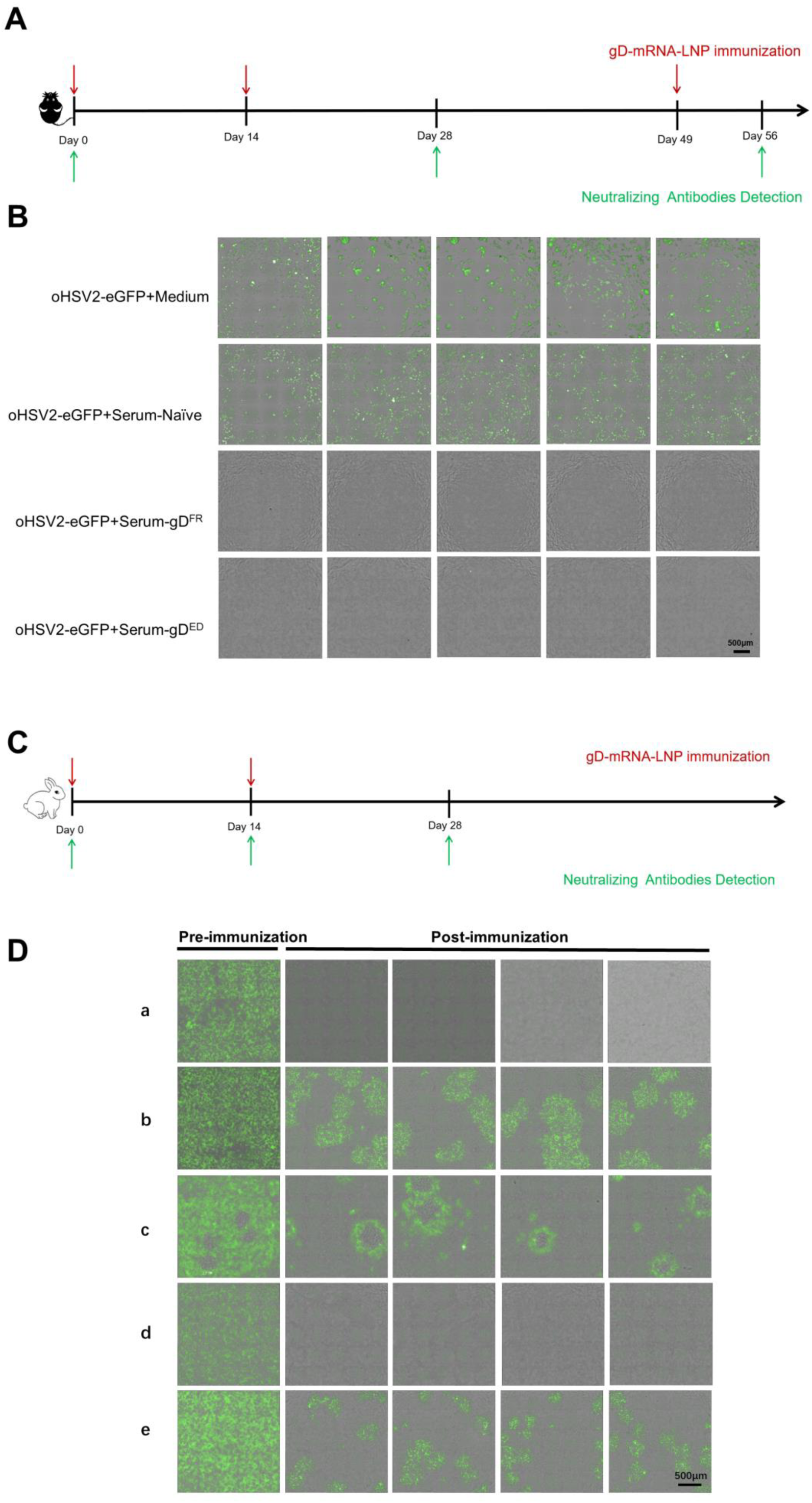
Detection of the neutralizing antibodies produced by animals immunized with gD^ED^-mRNA-LNP and gD^FR^-mRNA-LNP using Vero cells and virus-infected plaques. **A** Timeline of mouse immunization and blood collection. The red and green arrows represent the immunization time and the blood collection time, respectively. **B** The postimmunization serum (at day 56) neutralization of oHSV2-eGFP. DME/F-12 containing neonatal bovine serum (termed Medium), preimmunization serum (termed Serum-Naïve), postimmunization serum from a mouse immunized with gD^FR^-mRNA-LNP (termed Serum-gD^FR^) and postimmunization serum from a mouse immunized with gD^ED^-mRNA-LNP (termed Serum-gD^ED^) were incubated with oHSV2-eGFP. **C** Rabbit immunization regimen of gD^ED^-mRNA-LNP. Three rabbits received two doses of gD^ED^-mRNA-LNP immunization on days 0 and 14 (1 via intradermal injection, 2 via intramuscular). Blood was collected on days 0, 14 and 28 to separate the serum for the neutralization assay. **D** The serum neutralization antibody titers (on day 28) were determined using Vero cells and plaques observed 72 hours post-virus infection for: (a) the 20 μg intradermal injection group with serum diluted 64 times; (b) the 20 μg intradermal injection group with serum diluted 512 times; (c) the 25 μg intramuscular injection group with serum diluted 16 times; (d) the 50 μg intramuscular injection group with serum diluted 32 times; and (e) the 50 μg intramuscular injection group with serum diluted 64 times. The constant amount of virus was 100 CCID_50_ per well, and the serum (collected at 28 days) was diluted for neutralizing antibody detection.

In another experiment, rabbits were used to assess the immunogenicity of gD^ED^-mRNA-LNP in the induction of neutralizing antibodies. The initial immunization was designated as day 0. On days 0 and 14 of the experiment, 20 μg was injected intradermally into the hind neck of one rabbit, and 25 and 50 μg were injected into the muscles of the hind legs of the other two rabbits. At 0, 14, and 28 days, blood was collected from the middle ear artery to separate the serum to detect neutralizing antibodies (Figure 6C). To assess the neutralizing potency of antibodies produced by gD^ED^-mRNA-LNP after immunizing rabbits, live virus neutralization titrations were performed. Compared with the serum samples pre- and postimmunization, the serum samples postimmunization had clear neutralization activity of the virus (oHSV2-eGFP).

In intradermally injected rabbits, 1 μL of serum completely neutralized 40960 CCID_50_ viruses without eGFP expression or plaque formation (data not shown). Postimmunization serum from this rabbit had a neutralizing antibody titer (against 100 CCID_50_ viruses) of 512 (Figure 6D). However, the neutralization effect of the two rabbits injected intramuscularly was weaker than that of the intradermally injected rabbit. In the 25 μg intramuscularly injected rabbit, serum did not completely neutralize 100 CCID_50_ viruses when diluted 16-fold (Fig. 6D). In the 50 μg intramuscularly injected rabbit, when the serum was diluted 64-fold, the 100 CCID_50_ viruses could not be completely neutralized, and plaques appeared (Figure 6D). The neutralization titers for the immunized dose and route difference are shown in Figure S2. To exclude the effect of complement on the experiment, we performed complement inactivation. The experimental results showed that the effect of the complement inactivation group was the same as that of the non-inactivation group, and complement had no effect on the neutralization experiment (data not shown). It was concluded that rabbit serum samples following immunization (via intradermal or intramuscular injection) of gD^ED^-mRNA-LNP could neutralize the oHSV2-eGFP virus.

### 3.6. Cellular responses elicited in animals with uncapped gD-mRNA-LNP and S^δT^-mRNA-LNP

The ELISpot assay showed that animals immunized with gD^ED^-mRNA-LNP/gD^FR^-mRNA-LNP or S^δT^-mRNA-LNP produced specific T cells against the specific antigens. The primary immunization was recorded as day 0, and mice were immunized on designated days (Figure 7A, D). In the experiment for gD^ED^-mRNA-LNP/gD^FR^-mRNA-LNP, three groups (n = 8 per group) were designated: the gD^ED^-mRNA-LNP group (30 μg per animal); the gD^FR^-mRNA-LNP group (30 μg per animal); and the naïve group. On the 21st day, the spleens of 4 mice in each group were taken for ELISpot detection (data not shown), the remaining 4 mice in each group were given a second booster immunization with the same dose, and the spleens of those mice were taken on the 28th day to detect specific T cells. After mixing 3×10^5^ splenocytes with 10^6^ CCID_50_ of OH2 virus inactivated by UV (as stimuli) for 30 minutes and incubating for 48 hours, specific T cells were detected by the number of spots formed by IFN-γ cytokines secreted by splenocytes. As shown in Figure 7B, C for the data with spleen taken on day 28, the average number of spots formed by IFN-γ cytokines in the two groups (gD^ED^-mRNA-LNP and gD^FR^-mRNA-LNP) did have significant differences when compared with the average number of spots formed by IFN-γ cytokines in the naïve group. The results indicated that the two gD antigens translated *in vivo* by the uncapped mRNA structures could activate specific T cells in C57BL/6 mice. Similar results were obtained when mixing S^δT^-mRNA-LNP-immunized splenocytes taken on day 32 (Figure 7D) with SARS-CoV-2 S protein (as stimuli) followed by an ELISpot assay (Figure 7E and F).

**Figure 7.**
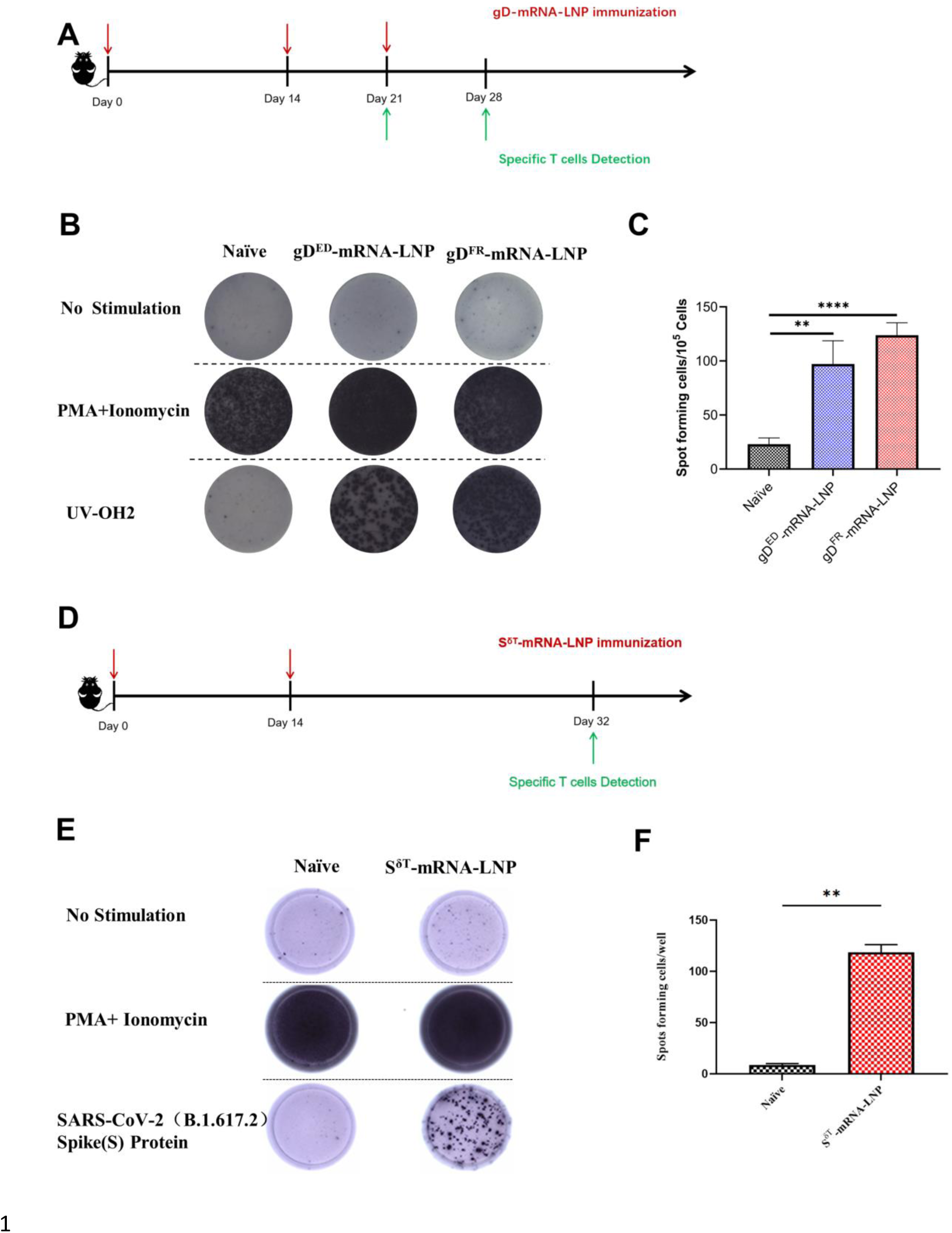
Animal T-cell immune responses after gD^ED^-mRNA-LNP/gD^FR^-mRNA-LNP or S^δT^-mRNA-LNP immunization. **A** and **D** Timelines of the gD-mRNA-LNP or S^δT^-mRNA-LNP immunization schedule and specific T-cell detection. The red and green arrows represent the immunization time and detection time, respectively. **B** and **E** The number of spots formed by IFN-γ cytokines secreted by splenocytes after immunization. **C** and **F** Quantification of spot formation by IFN-γ cytokines secreted by splenocytes after immunization. Student’s *t-*test (ns, not significant; *p < 0.05; **p < 0.01; ***p < 0.001; ****p < 0.0001).

### 3.7. Immunization with various mRNA-LNPs was able to induce high-titer specific antibodies

We also immunized rabbits with 4 different mRNA-LNPs encoding HSV2-UL19, HLA-E, NKG2A, and hGM-CSF. All these mRNA-LNP treatments induced high levels of specific IgG at 10^5^∼10^6^ titers. In addition, the antibodies were able to detect the antigens in their native forms: specific signals were obtained in immunofluorescence, flow cytometry, and western blot assays, in contrast to the pre-immune sera (Figure 8). These data suggest that mRNA-LNP immunization can be sufficient to prime animals and induce high production of activated B cells suitable to produce specific antibodies.

**Figure 8.**
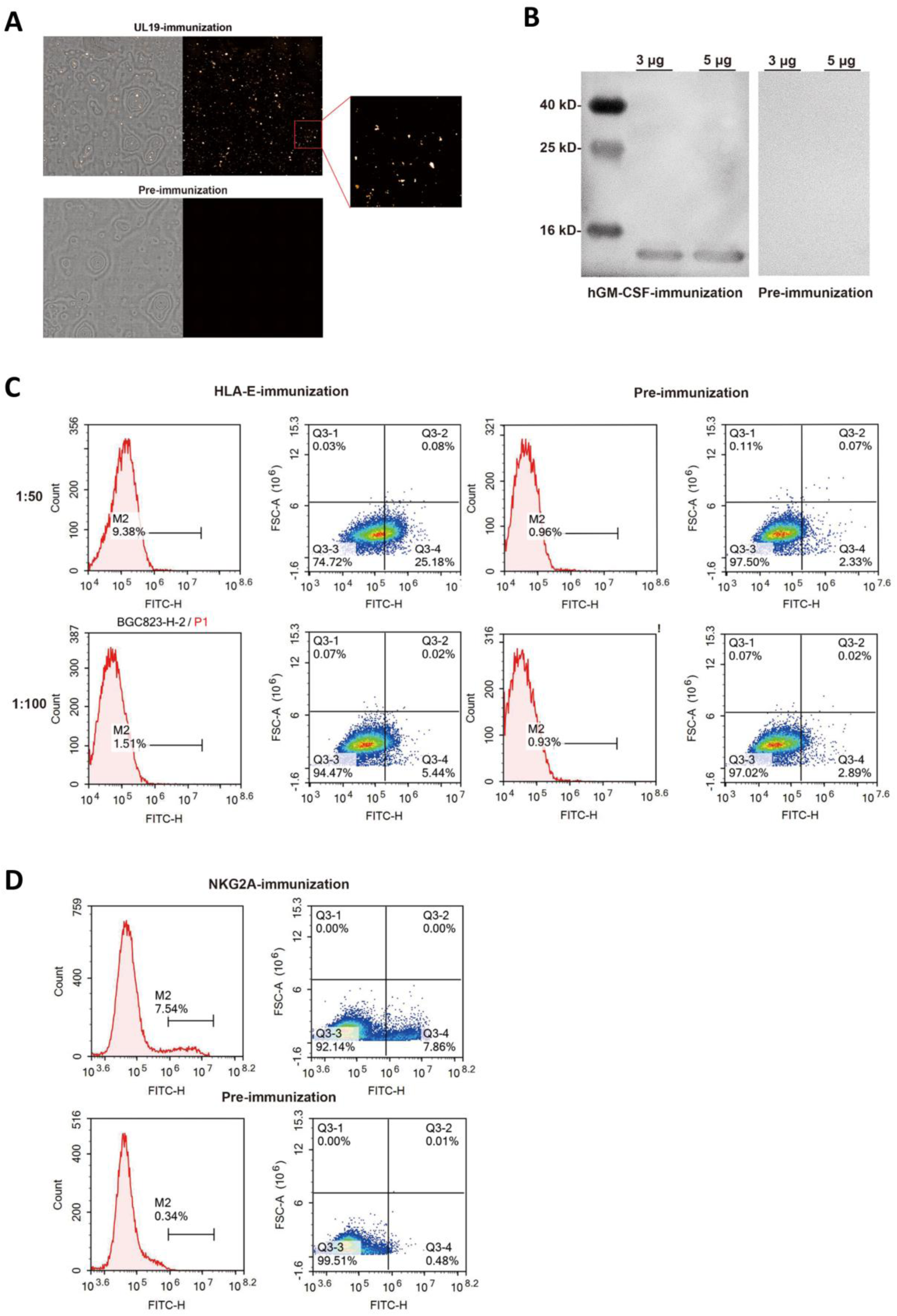
Analysis of the specificity of the rabbit antisera after mRNA-LNP injection. Rabbits were immunized 4 times with 50 μg of mRNA-LNPs by intradermal injection at 2 week intervals. Serum was collected 5 days after the final injection. **A** binding reactivity of HSV2-UL19-specific IgG levels to HSV2 particles was measured by immunofluorescence. The data were obtained at 1:1000 dilution. **B** IgG levels against hGM-CSF were determined by western blot. The data were obtained at 1:1000 dilution. **C** Serum from a rabbit injected with HLA-E mRNA-LNPs was tested for binding reactivity to BGC823 cells by flow cytometry assay. BGC823 cells were stimulated with HSV2 (MOI=1) to increase HLA-E expression. The antiserum of a rabbit immunized with HLA-E mRNA-LNPs displayed a positive reaction with BGC823 cells, while no signals were detected with the pre-immune serum. **D** NK92 cells was stimulated with IL-2 to increase NKG2A expression and incubated with serially diluted sera. The antiserum of a rabbit immunized with NKG2A mRNA-LNPs showed a rightward shift compared to its pre-immune control.

## 4. Discussion

Our mRNA platform has 4 notable features: Rapid, Amplified, Capless, and Economical (RACE; registered as the BH-RACE platform). Using IRES to replace the cap structure, using UTP but not N1MψTP to synthesize mRNA, and using self-developed encapsulation instruments and LNP delivery systems can greatly reduce the cost of raw materials, simplify the production process, and shorten the production time. At the same time, it has been shown that both the new coronavirus S protein mRNA vaccine and HSV2 gD protein mRNA vaccines can activate both humoral and cellular immunity in animals. Particularly exciting is that potent neutralizing antibodies against Delta and Omicron real viruses were induced with the new coronavirus S protein mRNA vaccine from the BH-RACE platform.

The application prospects of mRNA technology are very broad, and this technology can be used for the development of mRNA vaccines, protein replacement drugs and rapid antibody preparations. mRNA vaccines include prophylactic and therapeutic mRNA vaccines(To and Cho 2021). Prophylactic mRNA vaccines can express specific protein antigens when delivered into the body, thereby activating the immune response to prevent infectious diseases. Such vaccines are being widely developed for viruses including the new coronavirus (mRNA-1273, BNT162b2), the herpes simplex virus (Moderna, mRNA-1608; Binhui Bio, HSV2 gD mRNA vaccine)(Hassett et al. 2019), the influenza virus (mRNA-1010, BNT161, CVSQIV), the Zika virus (mRNA-1893), HIV (mRNA-1644), and the rabies virus (CureVac, CV7202, https://www.curevac.com/en/pipeline/).

Therapeutic mRNA vaccines are mainly antitumor vaccines. At present, a number of mRNA tumor vaccines are under development, mainly including Moderna pipelines (mRNA-4157, personalized cancer vaccine; mRNA-5671, KRAS vaccine), BioNTech pipelines (BNT111, advanced melanoma; BNT-113, HPV16+ head and neck cancer; BNT-122, adjuvant colorectal cancer), CureVac pipelines (CV-8102, oncology candidate; undisclosed, tumor-associated antigens), and Binhui pipelines (heterologous mRNA tumor vaccines). Regarding rapid antibody preparation, the mRNA-LNP antigen is designed to replace the traditional protein antigen, which facilitates the quick preparation of polyclonal and monoclonal antibodies. This fast antibody preparation technology has the advantages of many types of applicable antibodies, a short preparation period, high specificity and high purity. At present, our mRNA platform is used to prepare an HSV2 gD monoclonal antibody, which will be used to assist the development of an mRNA vaccine against HSV2.

The mRNA 5’ cap structure determines the “self” and “nonself” mRNA properties. Nonself mRNAs recognized by host cells can trigger strong host immune responses against nonself RNAs through toll-like receptors 3, 7-9 (Schlee and Hartmann 2016). Furthermore, the IRES sequence from Cytomegalo encephalitis virus used in our mRNA constructs could theoretically elicit a severe anti-RNA immune response leading to rapid RNA degradation and reduced protein expression over time, making it very unlikely to induce the antigen-specific immune response. However, our results showed that (1) *in vivo* transfection of our LNP-encapsulated Fluc mRNA (one of our uncapped mRNA constructs) resulted in high levels of expression with signal intensities as high as 10^8∼9^ total flux p/s, similar to the expression levels of *in vivo* transfection of LNP-encapsulated capped Fluc mRNA(August et al. 2021); (2) immunization of mice, hamsters and rabbits with our uncapped mRNA constructs induced high titers of IgG and neutralizing antibodies against the designed protein antigens, and (3) our uncapped mRNA constructs are able to induce target protein specific cellular immune responses.

mRNA vaccines activate humoral and cellular immunity in animals. The mice and rabbits were both immunized with our uncapped mRNA constructs encoding HSV2 truncated gD proteins. ELISpot (IFN-γ) assays (not performed on the rabbits) showed that the cellular immunity of the mice was activated, and the neutralization tests showed that sera from both animal species contained neutralizing antibodies. It was noted that immunization of rabbits via two routes (intramuscular and intradermal) resulted in the production of antibodies with different neutralizing titers, which is consistent with the previous report (Nicolas and Guy 2008). In addition, mice (C57BL/6) and Syrian hamsters were immunized with uncapped mRNA encoding the new coronavirus spike protein trimer of the Delta strain (S^δT^-mRNA-LNP). The ELISpot (IFN-γ) assay (not performed on the hamsters) showed that the cellular immunity of the mice was activated. ELISA could also detect serum IgG antibodies from both animal species. However, the serum IgG antibody titers of both animal species were significantly different. It has been reported that animals of different species immunized with the same mRNA vaccine could produce IgG antibodies with different titers (Vogel et al. 2021).

Our mRNA technology platform described here is notable for its uncapped mRNA construct and the omission of any nucleotide modification for the mRNA sequence. These characteristics form a clear contrast to those of the demonstrated and successful mRNA technology platforms developed by others, which are characterized by 5’ capped mRNA structures (Moderna, BioNTech and CureVac) and complete pseudouridine modification (Karikó et al. 2005; Pardi et al. 2018) (Moderna and BioNTech) on the mRNA sequences. Using neither the 5’ cap structure nor nucleotide modification undoubtedly not only reduces the cost of the raw materials but also simplifies the manufacturing process, representing a step forward in this field.

Although there is still a lack of sufficient studies about whether the IRES-driven translation of mRNA is affected by nucleoside modification in the primary RNA sequences, our data presented here suggest that nucleoside modification reduces antigen expression and the resulting antigenicity. We obtained a competitive high titer of total IgGs from our special mRNA platform. Another emerging and unanswered question is whether the total IgGs induced by the IRES-facilitated antigen expression from the unmodified mRNA are correlated with TLR recognition. Exogenous RNAs without nucleoside modification are treated as pathogens in mammalian cells and stimulate innate immunity via their interaction with TLRs, therefore leading to a short lifespan (Karikó et al. 2005) and theoretical insufficient target antigen expression levels. Our mRNAs containing the 5’ IRES structure without nucleoside modification successfully avoided elimination by the innate immune system. The mechanism underlying this phenomenon is interesting and needs to be explored further.

However, we should note that there still exist a few issues to be addressed. First, regarding the nucleoside modification in the mRNA sequences, we only compared the use of pseudouridine to replace the natural uridine but have not yet addressed other types of nucleoside modifications, such as m5C, m6A, N7-methylguanosine (m7G), or a combination of m5C and m7G. Last but not least, the uncapped mRNA constructs used in this work for the immunogenicity assays were mostly delivered using an LNP formulation developed by Moderna. Of note, a proprietary mRNA delivery system that might be more suitable for our proprietary mRNA structure is awaiting development.

## Acknowledgements

The study was supported by the National Major Scientific and Technological Special Project for “Significant New Drug Development” (2018ZX09733002), the Key Research and Development Program of Hubei province (2020BCA062).

## Availability of data and materials

All data generated or analyzed during the study are included in this article and/or the Supplementary Materials. Additional data related to the article could be requested on reasonable request.

## Ethical approval statement

All animal experiments were carried out in strict accordance with the requirements of the Hubei Provincial Committee for the Management and Use of Laboratory Animals and the ARRIVE Guidelines. Six- to eight-week-old female C57BL/6 mice and female BALB/c mice, 5-week-old female Syrian hamsters, and 5-week-old female Japanese white rabbits were purchased from the Hubei Provincial Laboratory Animal Research Center, Wuhan, China (License No: SCXK2020-0018). All animal experiments were approved by the Animal Ethics Committee of Hubei University of Technology (HBUT No. 2020016). Animals were housed in a controlled environment with a light/dark cycle of 12h and had free access to sterilized food and distilled water. The experiments were conducted after a period of one week adaptation.

