## Supplemental Figure S1 and S2 for "Development and application of an uncapped mRNA platform"

Figure S1. Neutralization antibodies detected for SARS-CoV-2 Delta and Omicron strains in four groups (0 µg, 25 µg, 50 µg, and 100 µg; n = 1 for each group).

Figure S2. The neutralizing antibody titers post gD^ED^-mRNA-LNP immunizations at various doses and via different routes.
